# From a single sequence to evolutionary trajectories: protein language models capture the evolutionary potential of SARS-CoV-2 protein sequences

**DOI:** 10.1101/2024.07.05.602129

**Authors:** Kieran D. Lamb, Joseph Hughes, Spyros Lytras, Francesca Young, Orges Koci, James Herzig, Simon C Lovell, Joe Grove, Ke Yuan, David L. Robertson

## Abstract

Protein language models (PLMs) capture features of protein three-dimensional structure from amino acid sequences alone, without requiring multiple sequence alignments (MSA). The concepts of grammar and semantics from natural language have the potential to capture functional properties of proteins. Here, we investigate how these representations enable assessment of variation due to mutation. Applied to SARS-CoV-2’s spike protein using in silico deep mutational scanning (DMS) we demonstrate the PLM, ESM-2, has learned the sequence context within which variation occurs, capturing evolutionary constraint. This recapitulates what conventionally requires MSA data to predict. Unlike other state-of-the-art methods which require protein structures or multiple sequences for training, we show what can be accomplished using an unmodified pretrained PLM. We demonstrate that the grammaticality and semantic scores represent novel metrics. Applied to SARS-CoV-2 variants across the pandemic we show that ESM-2 representations encode the evolutionary history between variants, as well as the distinct nature of variants of concern upon their emergence, associated with shifts in receptor binding and antigenicity. PLM likelihoods can also identify epistatic interactions among sites in the protein. Our results here affirm that PLMs are broadly useful for variant-effect prediction, including unobserved changes, and can be applied to understand novel viral pathogens with the potential to be applied to any protein sequence, pathogen or otherwise.

## Introduction

The conventional study of protein sequence variation requires the alignment of amino acid sequence data. Alignments help identify where variation accumulates within sequences, revealing the evolutionary constraints that dictate observed variation. In human genome studies, impactful missense mutations, non-synonymous substitutions resulting in amino acid changes, are primarily linked to disease, while in viruses, amino acid changes are more often linked to adaptive evolution^6,7^. Alternatively, these changes (and in fact the majority of possible missense mutations) are deleterious resulting in, e.g., misfolding and a decrease in fitness or are “neutral” and have little to no consequence.

To date approaches for assessing the impact of mutations are premised on the availability of comparative genome sequence data to assess features like the relative proportions of non-synonymous to synonymous substitutions (dN/dS) or evolutionary conservation at individual sites. Similarly, genome sequence-based surveillance methods usually require aligned real-time sequence data to assess the relative growth rate of a pathogen lineage^8,9^. Experiments on novel variants can be initiated when first detected, but they take considerable time to complete.

Natural language processing (NLP) methods can be applied to biological sequences like DNA and proteins^2,3,10^. Biological sequences are physical molecules represented by arbitrary characters. This is convenient since these characters, representing nucleotides or amino acids, can be processed in the same way as words in natural language. Hie. et al^2^ show how, much like natural language, biological sequence context can be used to create informative embeddings that reflect complex properties of the sequence. Specifically, concepts like grammar (the structural rules of a language) and semantics (the meaning of the language) can purportedly be applied to changes in protein amino acid sequences to capture ‘fitness’ and mutation effects, respectively^2^. The ‘grammaticality’ of a sequence is captured by its probability, indicating how well it conforms to the rules of protein formation and composition. The semantic difference between protein sequences can be measured by computing distances, a ‘semantic score’, between sequences within the embedding space.

Evolutionary scale model 2 (ESM-2)^3,10^ is a protein language model (PLM) trained on approximately 65 million unique protein sequences^3,10^. Here, we demonstrate that ESM-2 can infer fundamental properties that classically require a multiple sequence alignment. Applied to SARS-CoV-2 we show that the grammaticality and semantic score metrics reveal characteristic properties of the SARS-CoV-2 spike protein sequence (Figure 1). We further assess how to use these metrics for variant and sequence horizon scanning. Crucially, these metrics, produced from a single protein sequence using pre-trained models, bypassing the need for alignment data. We make a case for their future application in characterising proteins from emerging pathogens and assessing the constraints acting on protein sequences.

**Figure 1.**
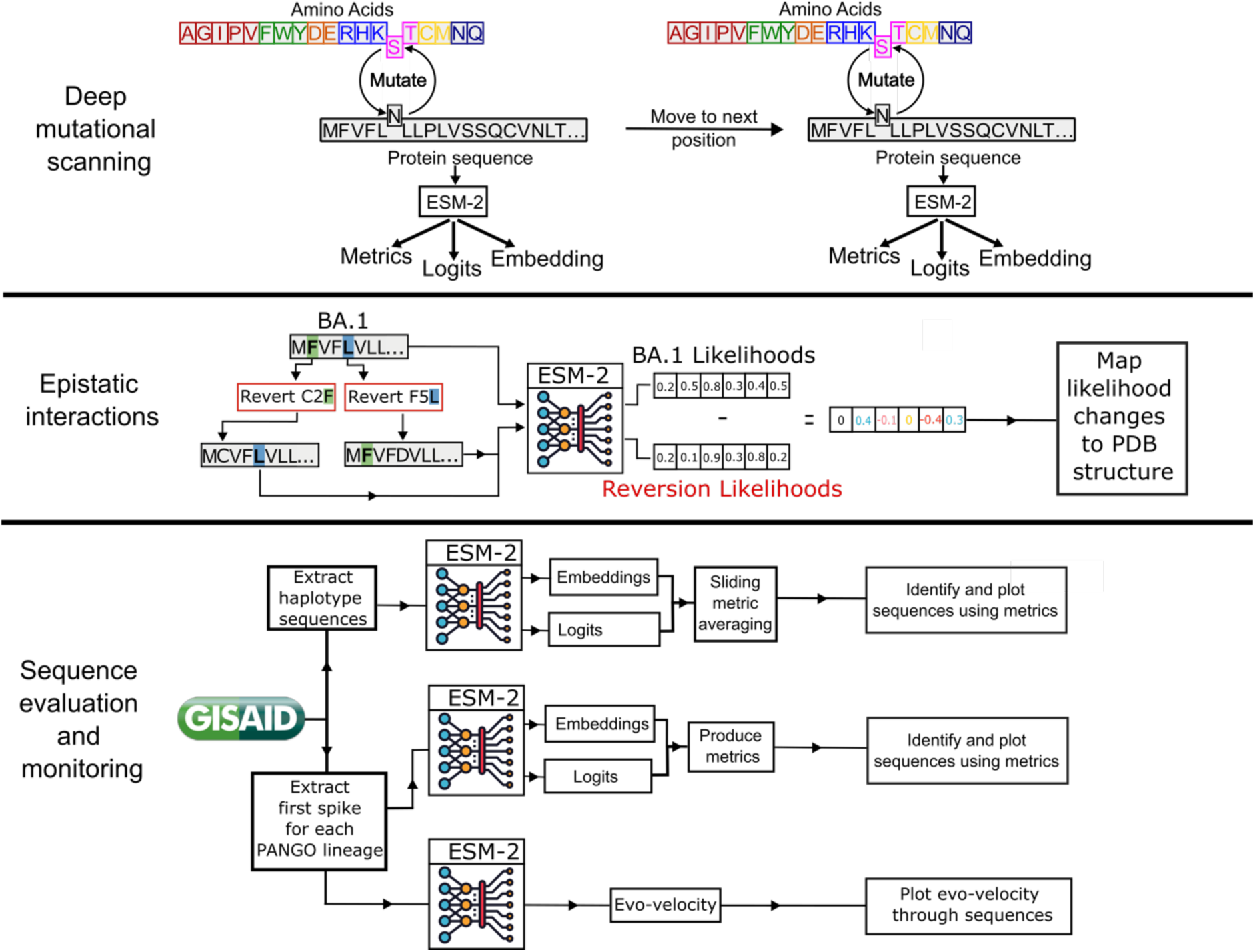
Schematic summary of our methodology. Deep mutational scanning involves taking a sequence and mutating every position to each of the possible alternative amino acids. These are then passed to the PLM to produce embeddings and logits which can be used for downstream tasks or to produce the metrics relative grammaticality and semantic score. Identifying epistatic interactions involves reverting the mutations from a SARS-CoV-2 variant (here BA.1) and measuring the effect this has on the other likelihoods. Embeddings and log-likelihoods (logits) can be used for surveillance by producing metrics over timescales for newly emergent protein sequences of concern, or by looking at evolutionary trajectories such as with evo-velocity.

## Results and discussion

### PLMs capture evolutionary potential

We first assess how well ESM-2 reflects the consequences of protein structure as a constraint on sequence variation. Specifically, does the model assign appropriate values for relative grammaticality (the log-likelihood ratio between the mutation and the reference amino acid) and semantic score (the distance between sequence embeddings, e.g., mutant and wild-type) at different sites in the protein sequence? To do this analysis, we utilised an in silico deep mutational scanning (DMS) approach^11^ using embeddings for every possible amino acid replacement in the SARS-CoV-2 spike protein. *In vitro* deep mutational scans involve experimentally substituting each site in a protein with every other amino acid and measuring the substitutions’ effect on phenotype. We do this computationally by embedding each of these mutated sequences using ESM-2 and then calculating grammaticality and semantic scores for each sequence.

The ESM-2 scores correlate well with the structural subunits of the spike (Figure 2). Using our DMS approach to produce relative grammaticality and semantic scores, we observe that the protein has two main regions that broadly correspond to the two subunits of spike protein S1 and S2 (Figure 2B and C). Relative grammaticalities decrease when substitutions are introduced in the S2 indicating that the region is less tolerant to change compared to S1. We observe a statistically significant difference between the mean relative grammaticalities of the S1 and S2 subunits (p-value = 2.168e-157, Mann-Whitney U test). Spike is a trimer with the core of the S2 subunit being formed by three closely interacting alpha helices from each monomer. As such, changes in this region could disrupt the formation of a stable and functional spike trimer by disrupting the inter-monomer interactions. On the other hand, in the S1 subunit, mutations in the N-terminal Domain (NTD) and receptor-binding domain (RBD) have higher relative grammaticalities. Despite being important to protein function, these regions need to be sufficiently flexible to facilitate receptor binding interactions and accommodate mutations to evade host immunity. This mirrors the accumulation of sequence variation in alignments. Specifically, using entropy, a measure of site-specific variation computed from multiple sequence alignments, there is significantly higher entropy observed at sites in S1 compared to S2 (Figure 2A), with each region found to be significantly different (p-value = 1.121e-10, Mann-Whitney U test). Thus, ESM-2 has correctly identified S1 as a more variable region than S2 (an analysis based on permutations of a single sequence) and indicates that ESM-2 has learnt this property about spike. The semantic rank also increases in the NTD region of the S1 (Figure 2A and B), indicating that changes here may produce large structural changes and are more likely to be accommodated due to higher relative grammaticalities across the region; presumably reflecting properties of the NTD, i.e., it is a region on the surface of the protein and contains many antigenic epitopes^12^. These results echo prior results that show PLMs can be used as an effective predictor of variant effect^13^. If ESM can model regional constraints, it may also be able to identify epistatic interactions that are linked with protein structural constraints.

**Figure 2.**
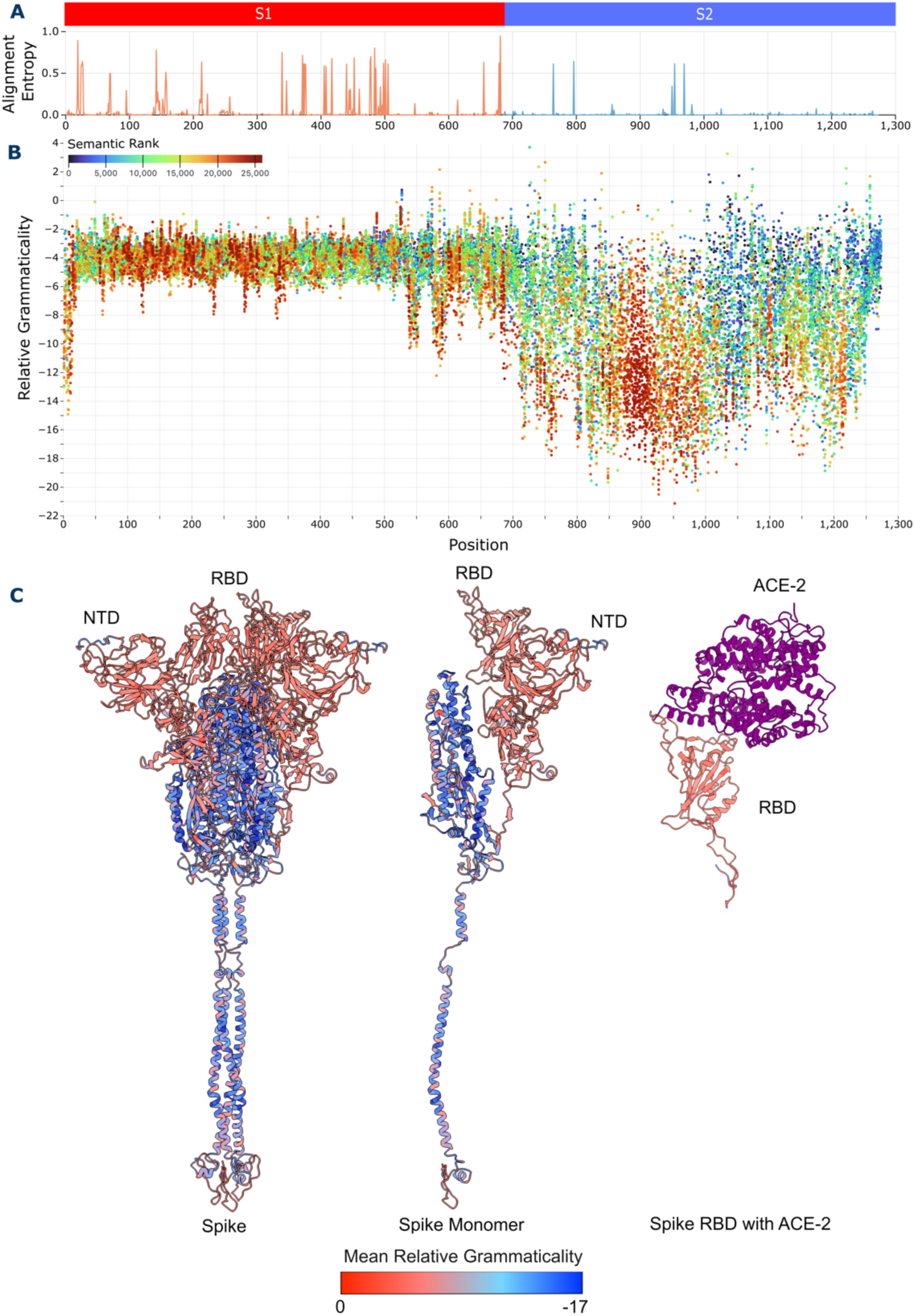
(A) Graph of the amino acid sequence alignment entropy at each position in the SARS-CoV-2 spike protein. S1 contains the majority of the sites with high entropy, while S2 contains only a few. (B) Scatter plot of spike protein DMS. Relative grammaticality is shown on the y-axis, with the amino acid position on the x-axis. Points are coloured on the semantic rank of each change. (C) Average relative grammaticality at each position on the spike protein plotted on three structures, the full Spike protein, the spike monomer, and the spike receptor binding domain (RBD) bound to the ACE-2 receptor in purple.

ESM-2 models spike protein constraints, maps low relative grammaticalities to the S2 and high relative grammaticalities to the S1, which is heavily targeted by host antibodies (NTD, RBD)^12,14–17^ and predominantly on the protein surface (Figure 2). Hie et al^2^ use the semantic score as a proxy for antigenic change, however, this is not necessarily relevant for protein sites that are not involved in antibody binding. Since ESM-2 is trained on a wide diversity of proteins, semantic scores represent shifts in structure/function rather than necessarily antigenic change. Many high semantically ranked changes occur in the NTD (Figure 2B) which contains a “supersite” with several epitopes targeted by neutralising antibodies^12^. The NTDs low relative grammaticality and high semantic score suggests it allows large structural changes to alter its antigenic properties without greatly affecting function.

### PLMs identify epistatic interaction sites

A consequence of protein structural constraint is that compensatory mutations are often required before a change can be accommodated. Such epistatic effects can be due to direct or indirect mechanisms linked to residues in close contact in the protein structure conformation and/or protein stability^18^. Here we present a novel approach for assessing epistasis, where we find putative epistatic interactions among sites by identifying where changes in the amino acid at one site cause significant changes in amino acid likelihoods at another. Language models compute likelihoods for each amino acid in a sequence based on the context of the rest of the sequence. This means that the likelihoods depend on the other amino acids in the sequence and will change if other positions mutate. Positions that are unaffected by the mutated site would be expected to change minimally, whereas those affected may change their probabilities, i.e., lower or higher depending on whether an epistatic interaction is antagonistic or synergistic, respectively. For example, the BA.1 Omicron, variant of concern (VOC) is defined by over 30 unique substitutions in its spike protein sequence. By reverting these defining mutations to the Wuhan-Hu-1 reference sequence amino acid, we measure the effect this reversion has on the probabilities of all other BA.1 amino acids computed by the model. To illustrate these results we highlight the interactions of three key substitutions E484A, D614G and N969K^19–22^; selected due to their effects on antigenicity, conformational changes and presence in the S2 domain:

The E484A substitution in the RBD of spike is involved in modulating binding to the ACE-2 receptor and is a good example of a substitution with localised effect (Figure 3A). The VOCs Beta and Gamma contained the E484K substitution and this contributed to escape from several RBD-neutralising antibodies^23,24^. Interestingly, A, D, G and K substitutions at this site 484 all confer resistance to human convalescent sera^16,23^. E484A shows decreased immune evasiveness relative to E484K, however interactions between N501Y and Q498R mutations recover this^20^. Reversion of the mutation changes amino acid likelihoods primarily within the RBD (Figure 3A).

**Figure 3.**
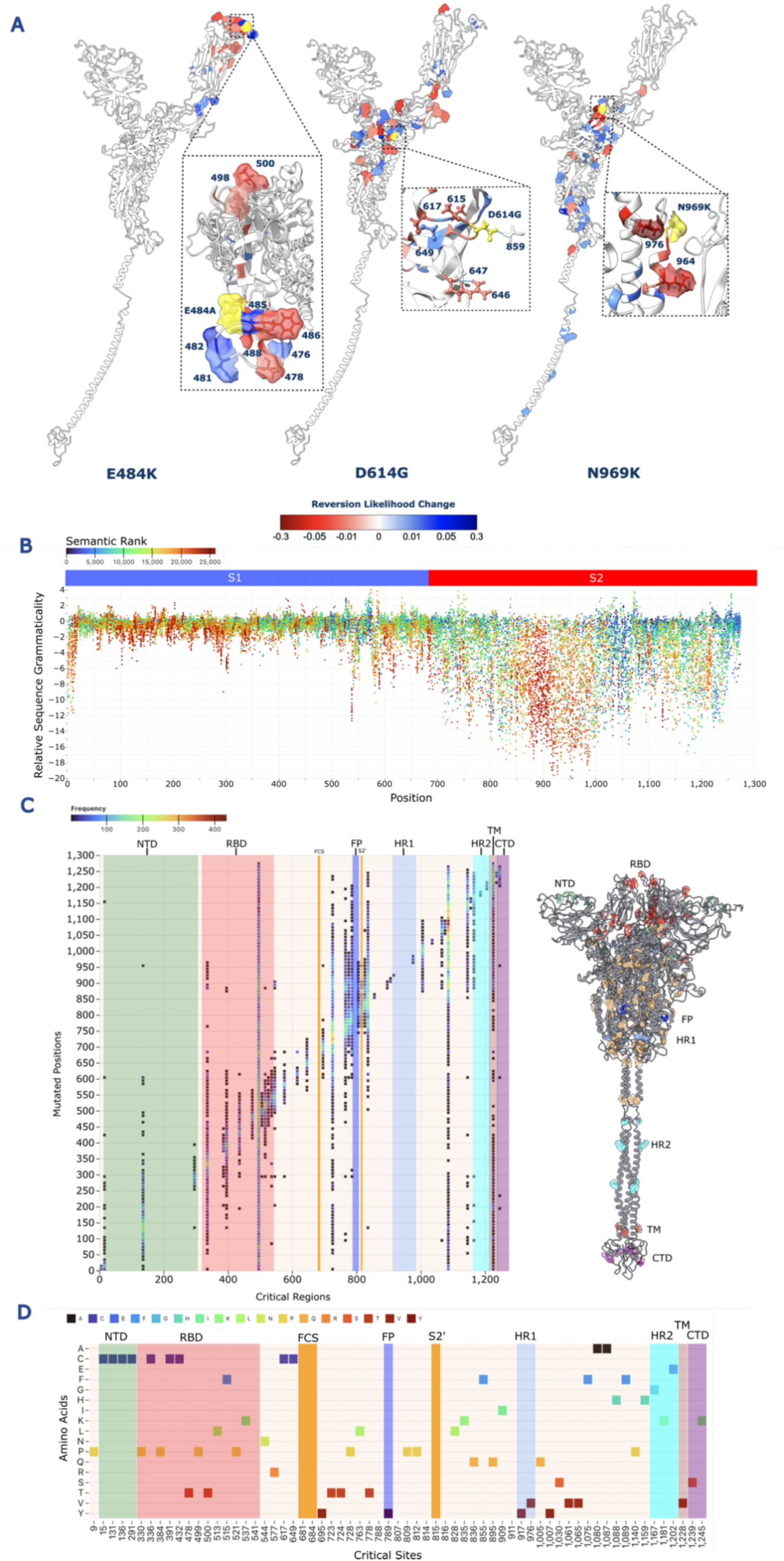
(A) Monomeric structures showing the changes in probabilities for three example substitutions in SARS-CoV-2’s spike protein: E484A, D614G and N969K. The mutation site is coloured yellow, blue sites increase in probability while red sites decrease. Mutation probabilities were only shown if they were outside two standard deviations of the mean change. (B) Relative sequence grammaticalities, the product of each amino acid likelihood rather than just the mutations, against the amino acid position. Amino acids are coloured on the semantic rank, which is a ranking of the semantic scores of all positions from highest to lowest semantic score. (C) The significantly changed logits across the whole DMS were identified, with positions that were repeatedly identified (called critical sites) as being affected by the in-silico mutations counted. These were then mapped onto the spike structure and coloured on their domains. (D) Amino acids of each consistently affected reference residues from the DMS data. The NTD and RBD contain mostly Prolines(P) and Cystines(C), while the rest of spike has a wider distribution of amino acids.

RBD positions 485, 486 and 488 have the largest absolute changes in probability. Site 485 increases its likelihood and thus appears to be antagonistic with E484A, indicating incompatibility with the mutation. Positions 486, 488 and Q498R reduce their likelihoods upon E484As reversion, indicating that the mutation is synergistic with these positions, possibly improving the structure or at least minimising disruption.

Our results show S1 reversions have shorter distances to their affected positions in sequence space and in three-dimensional space (Supplementary Figure 3) likely due to it being the subunit comprising most of the accessible surfaces of the protein. Changes more ‘central’ to the protein structure like those in the S2 domain have a greater chance to affect more amino acids due to being almost completely surrounded by other sites. This can produce more impactful changes that have knock-on effects at distant locations. While 501Y is not identified, position 500 decreases its likelihood further suggesting synergistic interactions with this region.

The D614G substitution emerged early in the pandemic^25^, is now present in all circulating lineages and is an example of a substitution with broad effect (Figure 3A). Specifically, D614G has two main functional consequences: (1) to remove a hydrogen bond between 614 and 859 on the adjacent spike monomer, and (2) at 647 to contribute to the structure of the C-terminal domain. 614G increases the stability of the spike protein^19^ as well as the infectivity of the virus at the cost of increased susceptibility to neutralising antibodies^19,26^. Its reversion affected 45 other positions (Supplementary Figure 4), nine of which are in the RBD, with a further 30 in S1 and 6 in S2. While 859 and 647 were not affected positions, 615, 617 and 649 did change their likelihoods. 649 is an internal amino acid located behind 614 while 615 is in direct contact with the site and 617 is slightly downstream. 617 and 615 both reduced their likelihoods, indicating that the D614G acquisition was synergistic with those sites. 649 increase its likelihood by 3.7% on reversion suggesting an antagonistic effect with 614G.

The N969K reversion affected 46 positions on the protein (Figure 3A, Supplementary Figure 3 and Supplementary Figure 4) and has been linked to the change in Omicron entry route^21^. N969K forms electrostatic contacts with Q755 on the adjacent protomer in the pre-fusion state^27^, as well as interacting with and displacing the HR2 backbone in the post-fusion spike structure^22^. It has long-range interactions both in linear sequence and protein structure context (Supplementary Figure 3A) and has the greatest number of affected sites compared with the other BA.1 mutations. Position 76 had a likelihood drop of 21% while position 964 had a 15.9% decrease, indicating a strong synergistic effect with the N969K substitution. N969K is positioned in the middle of a loop connecting two alpha helices, with 976 and 964 positioned at the end of the first helix and the start of the second. The large likelihood shift indicates strong synergy with the N969K mutation and could be as yet undiscovered epistatic positions. Both D614G and N969K had positions of interaction on other monomers, however ESM was only given a single monomer, so the lack of other monomers in the embedding may prevent the detection of these interactions.

Position 330 is of particular note in our analysis as it decreases its likelihood in the three reversions we characterised (E484A, D614G and N969K), but also in several other mutations in BA.1. This prompted an investigation into whether certain sites were more prone to fluctuations than others, and whether these sites were of functional importance. We identified positions that significantly changed their likelihoods in each DMS mutant, before filtering for sites that occur across many different mutations and investigating their functional relevance to the protein. Figure 3C shows several sites in the protein that consistently change their likelihoods in response to changes elsewhere. Strikingly, positions 499, 723, 1087 appear to be affected by substitutions across the whole protein while other positions such as 330 tend to have effects at closer ranges. The amino acids in the NTD and RBD that consistently change their likelihoods are mostly prolines and cysteines, while elsewhere there is a larger spectrum of amino acid types, eight of which contain aromatic side chains (F and Y) (Figure 3D). These amino acids have important structural features. Proline is a rigid amino acid due to its side-chain which reduces its flexibility^28,29^ and prevents it from forming stable a-helices^28^. Prolines are often found where sharp turns are necessary for structure. Cysteine thiol side-chain allows them to form covalent di-sulphide bridges between adjacent cysteines^30^. Cysteines are well conserved throughout proteins^30^, and several of the identified cysteines were involved in di-sulphide bridges (Figure 3C and Figure 3D).

Since likelihood changes across the protein identify epistatic and structural effects, we can produce whole sequence grammaticalities that measure the effect of a mutation on the whole of the sequence (Figure 3A and B). This relative sequence grammaticality is the product of the mutated sequence logits (a pseudo-log likelihood)^31^. While some mutations are deemed to be unlikely by the PLM, selection pressure may allow these changes to persist if they are beneficial. We see in Figure 3D a similar distribution to our DMS results (Figure 2B), however many more positions are now positive indicating that the epistatic effect of the mutation has made the sequence more likely than the reference. Several important substitutions that have been observed in VOCs are among these including D614G, E484K/A, K417N, P681R and more. Each of these mutations make the sequence less likely compared to Wuhan-Hu-1 when measured with relative grammaticality. D614G has a negative relative grammaticality of −3.07, yet its sequence grammaticality is positive at 0.105, indicating that the overall sequence is fitter with 614G than 614D based on the model. E484A has a positive relative sequence grammaticality of 0.151, while the mutation has a negative grammaticality of −2.53. N969K remains negative (−11.31 relative grammaticality, −2.90 relative sequence grammaticality), indicating that it still appears to reduce sequence fitness. This presumably represents a fitness trade off as it appears N969K destablises the proteins post-fusion conformation^22^, while stabilising it pre-fusion context^27^.

### PLMs provide new metrics for assessing mutational impact

An important question is the relationship between the PLM metrics and conventional experimental, structural or evolutionary metrics, or does grammaticality and semantic score represent a novel measure of change? Extensive high-throughput experimental investigations have revealed various aspects of spike biology. We sought to correlate these measurements with PLM metrics, to better understand what characteristics are captured by the model. We calculated spearman’s rank correlations between the embedding metrics and experimental and computational metrics. We then used the full embeddings and logits as well as score combinations to fit linear and support vector regressions (SVR) to each metric and again scored using a spearman’s rank correlation.

Experimental scores were used from three *in vitro* DMS studies, two of which were performed on the RBD while the other was performed on a full spike protein. “*Escape*”, “*Entry*” and “*Binding*” were determined from a full spike DMS by Dadonaite et al. (2023)^32^ and correspond to: immune escape from human sera, cell entry, and binding affinity to soluble ACE-2 respectively. “*RBD Escape*” was produced by Yisimayi *et al.* (2024)^33^ using an RBD only DMS. The “*Wuhan*” and “*Variant*” binding and expression scores are from an RBD only DMS from Starr *et al.* (2022)^34^ where binding to ACE-2 and expression of the protein were measured across five spikes. The “*Variant*” score is made from an average of both measurements across all five spikes.

Interestingly, neither grammaticality nor the semantic score correlated with the full spike experimental DMS scores; with coefficients below 0.2 for “*Binding*” and “*Escape*”. “*Entry*” was better correlated with the PLM scores, with correlation coefficients greater than +0.3 and less than −0.3 for grammaticality and semantic score respectively. “*Escape*” correlation coefficients were also below 0.2 for all metrics both for the full spike and the RBD. The semantic score showed stronger negative correlations with RBD based DMS results of around −0.3 (Figure 4A). Fitted linear regressions improve the correlation coefficient for the *“RBD Escape”* with the mean embedding and the Wuhan-Hu-1 *“RBD Binding”* and *“RBD Expression”* when using sequence logits (Figure 4B). However, linear models failed to improve much upon metric correlations likely due to their inability to exploit non-linear relationships in the high-dimensional representations. SVR improves substantially here, with a correlation coefficient between 0.58 and 0.73 for all experimental measurements aside from “*Escape*” and *“Binding”*, although both improve their coefficients over the metric and linear model correlations (Figure 4C).

**Figure 4.**
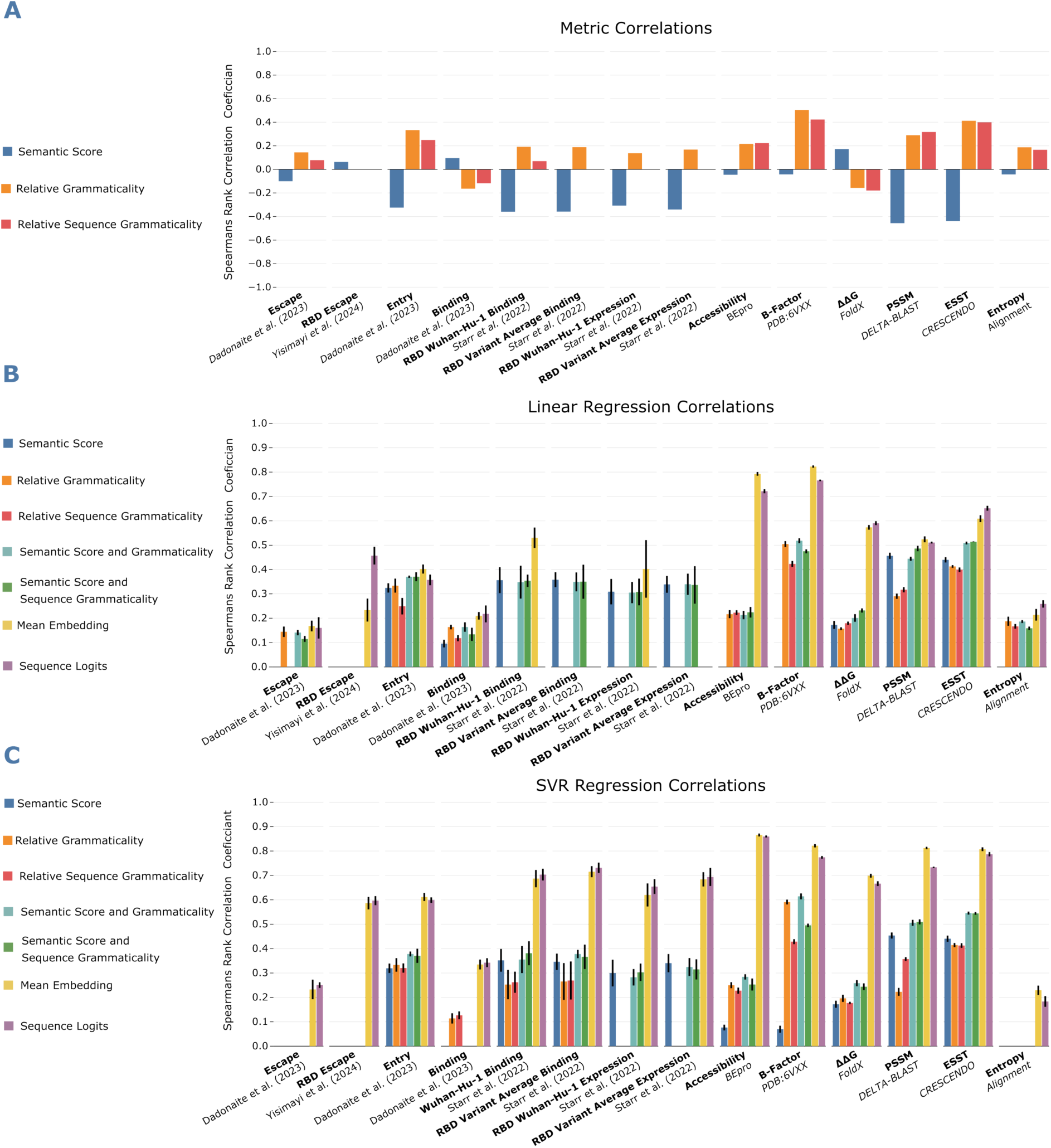
(A) Spearman’s Rank correlations between the semantic score, grammaticality and relative sequence grammaticality and traditional metrics. Bars not present in a metric category means the correlation was not found to be significant after a Bonferroni correction. (B) Spearman’s Rank correlations between the language model metric and the traditional metric. Each pair was fitted using a grid search and a linear regression model, with 5-fold cross validation. Bars represent the mean of the correlations, with the error bar +/- 1 standard deviation of the correlations. Bars not present in a metric category means the correlation was not found to be significant after a Bonferroni correction. (C) Spearman’s Rank correlations with a support vector regression model using an RBF kernel and 5-fold cross validation.

We also calculated several computational metrics that are commonly used for the analysis of protein sequences and estimation of evolutionary constraint. These provide quantitative measures of substitution probability and structural characteristics at every site in the protein and are intended to be representative of the wide range of techniques currently used to analyse protein structural evolution. This allowed for comparison of the PLM metrics with existing metrics.

The crystal structure of the spike protein was used to calculate “*Accessibility”,* “*B-Factor”,* “*ΔΔG*” and the environment-specific substitution table (ESST). “*Accessibility”* measures the antibody accessibility at every position. The “*B-Factor”* is the temperature factor derived from the protein crystallography experiment used to determine the 6VXX structure and is a measure of the local fit of the structure to experimental data; it is often increased if there is static or dynamic disorder. Protein stability change (ΔΔG) was assessed with the FoldX software. The ESSTs were taken from Mizuguchi et al.^35^ and provides a statistical estimate of substitution likelihood based on the observed frequency of substitutions at amino acids in similar local amino acid structural environments. We also calculated two sequence-based measure of substitution likelihood. The position-specific scoring matrix (PSSM) was calculated using DeltaBlast and calculates the log likelihood of substitutions occurring based on their frequency in related sequences^36^. “*Entropy”* was used to provide an estimate of the variability present at a site.

The semantic score showed strong negative correlations coefficients of <-0.4 with the computational likelihood metrics *“ESST”* and “*PSSM”*. Grammaticality scores performed similarly or slightly worse for the likelihood metrics although the scores were inversely correlated compared to the semantic scores. *“B-Factor*” had the best correlation with relative grammaticality and relative sequence grammaticality, with both having correlation coefficients greater than 0.4. This indicates that grammaticalities (which align well with conservation) may also be a reasonable proxy for assessing protein flexibility.

Both linear regression and SVR drastically improve correlations for the computational metrics. The linear model produced correlation coefficients >0.5 for all measurements except “*Entropy”*, although this might be expected since it’s a site-specific metric rather than mutation specific like DMS. “*Accessibility”* and “*B-Factor”* with both logits and embeddings achieve high correlation coefficients of >0.7 while *“ΔΔG”*, also improves with a co-efficient of >0.5 for logits and embedding features. SVR improves all correlation coefficients across the board except for “*Entropy”*. *“PSSM”* experiences the greatest increase from ~0.5 to ~0.8 using logits and SVR.

We observe that the PLM scores (semantic score, relative grammaticality and relative sequence grammaticality) correlate only weakly with the tested experimentally or computationally derived metrics. However, at least one of the language model metrics significantly correlated with all the other metrics (Figure 4A). These experiments show that embedding metrics are not simply an alternative way to describe existing metrics.

When a significant correlation was achieved, linear regression predictions for logits and embeddings either match or drastically outperform correlations using the model metrics, or their combinations (Figure 4B). Fitting an SVR results in more consistent and higher correlation coefficients across the board, with only *“Binding*”, *“Escape”* and *“Entropy”* achieving correlations of less than 0.5 with logits and embeddings as features (Figure 4C). Score combinations show little improvement over individual scores.

We also calculated a statistical distance metric, the Jensen Shannon distance (JSD), between these computational metrics and the semantic score and grammaticality at every site. Assessing the mean JSDs across different regions of the spike protein reveals a statistically significant difference between regions in the S1 and the S2 for the grammaticality score vs other metrics (Supplementary Figure 6). This supports a conclusion that grammaticality carries implicit context-aware information about differences in evolutionary constraint between protein regions, as indicated by Figure 2B; similar information not captured by existing, site-independent computational metrics. JSD between semantic score and the other metrics does not show pronounced differences between S1 regions and S2 as consistently, although there were still statistically significant differences between some of the protein regions, again demonstrating contextual structural awareness in the PLM embeddings.

### Language models recapitulate evolutionary relationships between related sequences

The earliest known SARS-CoV-2 sequence for each PANGO lineage was extracted from the global data (retrieved from GISAID) and their spike protein sequences embedded using ESM-2^3,10^. The evo-velocity package was then used to produce an evo-velocity UMAP for the sequence embeddings. Evo-velocity assigns a putative directionality between the embeddings that describes the flow of evolution through the UMAP embedding space. Firstly, evo-velocity^37^ shows that PLM embeddings can distinguish between meaningfully different spike proteins (Figure 5A and B). Secondly, it illustrates that embeddings can help to understand the evolutionary landscape of spike and the directionality of its evolution.

**Figure 5.**
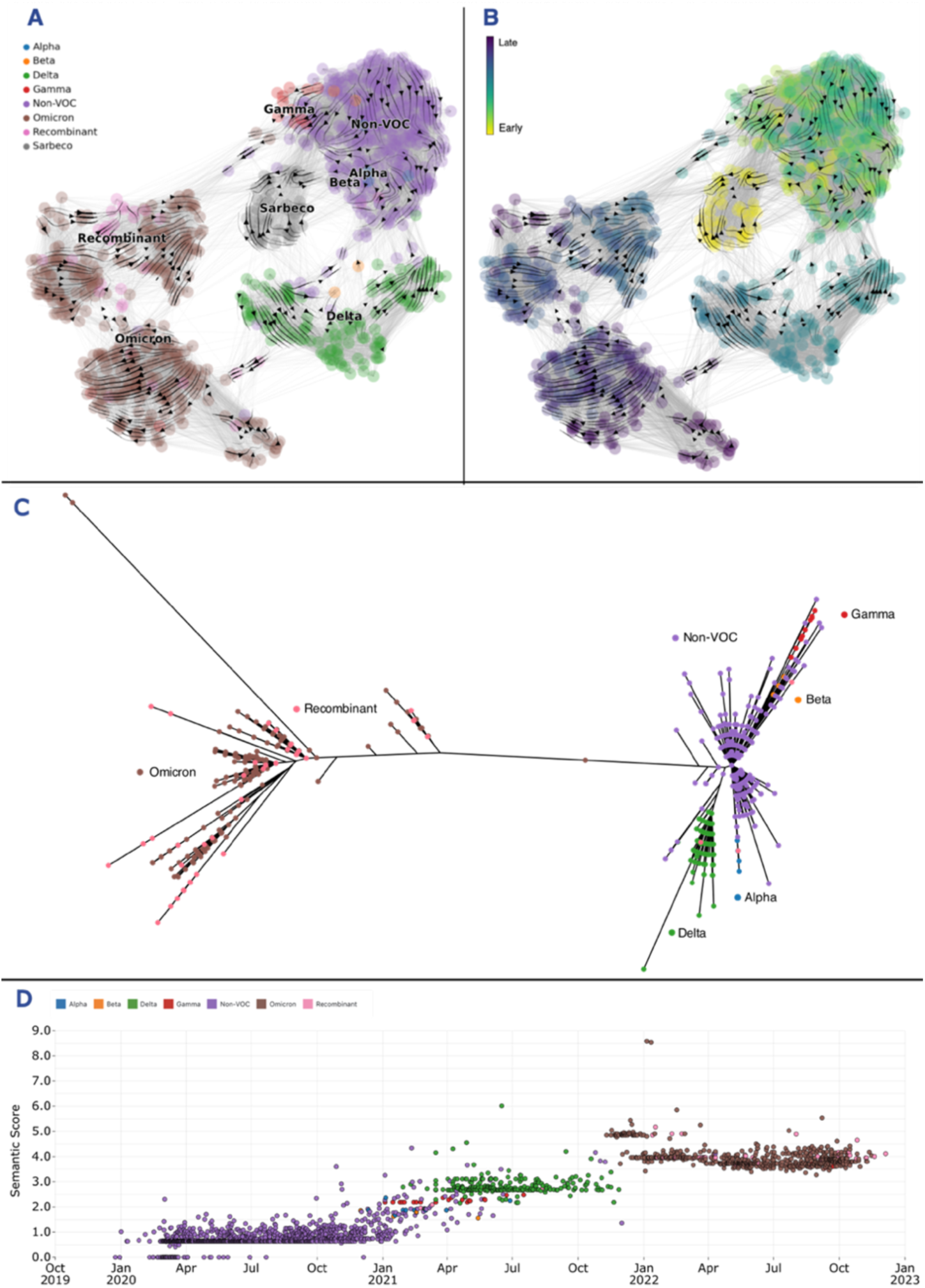
(A) UMAP of initial spike sequence embeddings for SARS-CoV-2 PANGO lineages and a selection of other known Sarbecovirus spike sequences. Each lineage is represented by one spike embedding. Points are coloured on VOC classification. Arrows represent the evo-velocity through the embedding space, which shows a “directionality” of evolution. (B) shows the sequences coloured by pseudotime inferred using sequence embedding probabilities to order sequences in time using an inferred root and an endpoint. (C) Shows an unrooted nucleotide phylogenetic tree of the spike sequences, coloured again by VOC. (D) shows the spike protein sequences plotted using their sample date and semantic score coloured consistently to Figure 1B.

The evo-velocity accurately describes the evolution of SARS-CoV-2 moving from the earlier ‘non-VOC’ clusters into VOC clusters, consistent with their evolution during the pandemic (Figure 5A and B). Omicron and Delta form distinct clusters while Gamma forms a concentrated cluster on the fringes of the non-VOC sequences. Beta and Alpha are less homogenous, and recombinants tend to fall with their parental lineages (primarily Omicrons). Sarbecoviruses (which include bat and pangolin viruses related to SARS-CoV-2) form another distinct cluster close to the early SARS-CoV-2 sequences. These are also identified as the earliest sequences by the evo-velocity pseudotime algorithm (Figure 5B and Supplementary Figure 5). Evo-velocity first uses diffusion analysis to identify root and endpoint sequences before estimating the order of evolution using a pseudotime simulation (Figure 5B and Supplementary Figure 5). Pseudotime analysis achieved a significant (p-value = 3.24e-296) spearman’s rank correlation of 0.86 against sampling time, confirming that the model has inferred the evolution of the sequences in the correct order. Evo-velocity recapitulates the topology of the representative sequence phylogeny in Figure 5C, with early VOCs more closely related to non-VOC sequences than Omicrons and recombinants. This shows PLM embeddings capture meaningful representations that differentiate between distinct sequences. The congruence between the phylogenetic tree topology and the evo-velocity derived structure provides further confirmation that this method captures the evolutionary history of the spike protein.

PLM metrics also differentiate between distinct SARS-CoV-2 spike proteins. Using the semantic score or the relative grammaticality, a non-VOC, early VOC (Alpha, Beta, Gamma and Delta), and an Omicron cluster can be identified (Figure 5, Supplementary Figure 7 and Supplementary Figure 8). Unlike the UMAP (Figure 5A and Supplementary Figure 5) which is a dimensionality reduction and projection from high-dimensional space, grammaticality and the semantic score are much simpler to produce. Despite this, they still recapitulate much of the information shown in the UMAP.

Evo-velocity, thus, captures differences between VOC sequences, recapitulates the topology of the nucleotide sequence phylogeny and reconstructs the direction of evolution accurately (Figure 5). The PLM scores can effectively group lineages into their VOC categories (Figure 5D, Supplementary Figure 7 and Supplementary Figure 8).

### Detecting the distinct nature of the variants of concern on emergence

The emergence of SARS-CoV-2 and its subsequent variants of concern (VOCs) caused large waves of infections during the pandemic. Predicting their fitness advantage just from their initial sequences, before the viruses were in widespread circulation, has proven incredibly difficult. PLMs could be used to meet this challenge of rapid characterisation of individual pathogen genomes as we have implemented for relative grammaticality and semantic scores, see the ‘Variant Assessment’ tab at https://sars2.cvr.gla.ac.uk/cog-uk.

The PLM metrics produced for each PANGO lineage can be used to assess SARS-CoV-2 variants since they detect differences between sequences. We observed a characteristic “jump” in both semantic score and relative grammaticality between non-VOC, early VOCs and Omicron sequence clusters (Figure 5D and Supplementary Figure 8). Earlier VOCs may have required fewer changes to compete since most of the human population was still naive to infection or vaccine-derived immunity. The Omicron lineage emerged during high levels of both vaccination and infection. It contained many more substitutions in the spike protein than previous variants, it changed its entry mechanism preference and managed to largely evade previous immunity which resulted in an extended vaccination regimen of three doses being recommended^38,39^. Unlike the other VOC groups, Omicron sequences initially decreased their semantic scores. The BA.2 variant had fewer mutations than the initial BA.1 variant, however later sequences that increased their number of mutations, continued to decrease their semantic scores. Clearly the semantic score is not a proxy for mutation count.

Once regular sequencing and surveillance are underway, a pipeline for analysing emerging sequences could identify outliers that may become future VOCs. Despite PLM scores distinguishing between VOCs well, a larger score does not always mean a variant will be successful. This could be due to measuring against Wuhan-Hu-1, which is now an extinct and increasingly irrelevant SARS-CoV-2 variant. To tackle this, we created a dynamic embedding that represents the average sequence circulating at a given time. This means we can check whether sequences are both divergent from the SARS-CoV-2 reference or from what is currently or recently circulating. By looking at a UK subset of sequences, we can assess this approach thanks to the UKs high sequencing capacity paired with well-defined lineage waves. UK sequences were clustered to 99.9% similarity and a representative haplotype sequence for each cluster was embedded using the model.

Waves of semantic change take place between all major VOC waves in the UK (Alpha to Delta to BA.1 to BA.2) (Figure 6). The non-VOC to Alpha transition resulted in sequences deviating by semantic score of >2, although this quickly increased to almost three after the emergence of Delta. Following BA.2, the semantic changes in Omicron have been more incremental, with a gradual increase up to March 2023, followed by a decrease until July 2023. There have been several smaller waves after Omicron’s emergence, which suggests repeated dominance and replacement of consecutive distinct sub-lineages throughout this period. The haplotype embeddings also uncover the large diversity of semantic scores contained outside just the PANGO representative sequences from Figure 5D. Alpha and Delta have a large range of semantic scores, equalling some of the most divergent Omicron sequences despite pre-dating them by months (Figure 6C). To a lesser extent, the non-VOC sequences also have some very high semantic scores relative to the average semantic score during the Delta wave. What is clear is that as a new VOC began to take over, sequences diverge very quickly from the average circulating sequence embedding.

**Figure 6.**
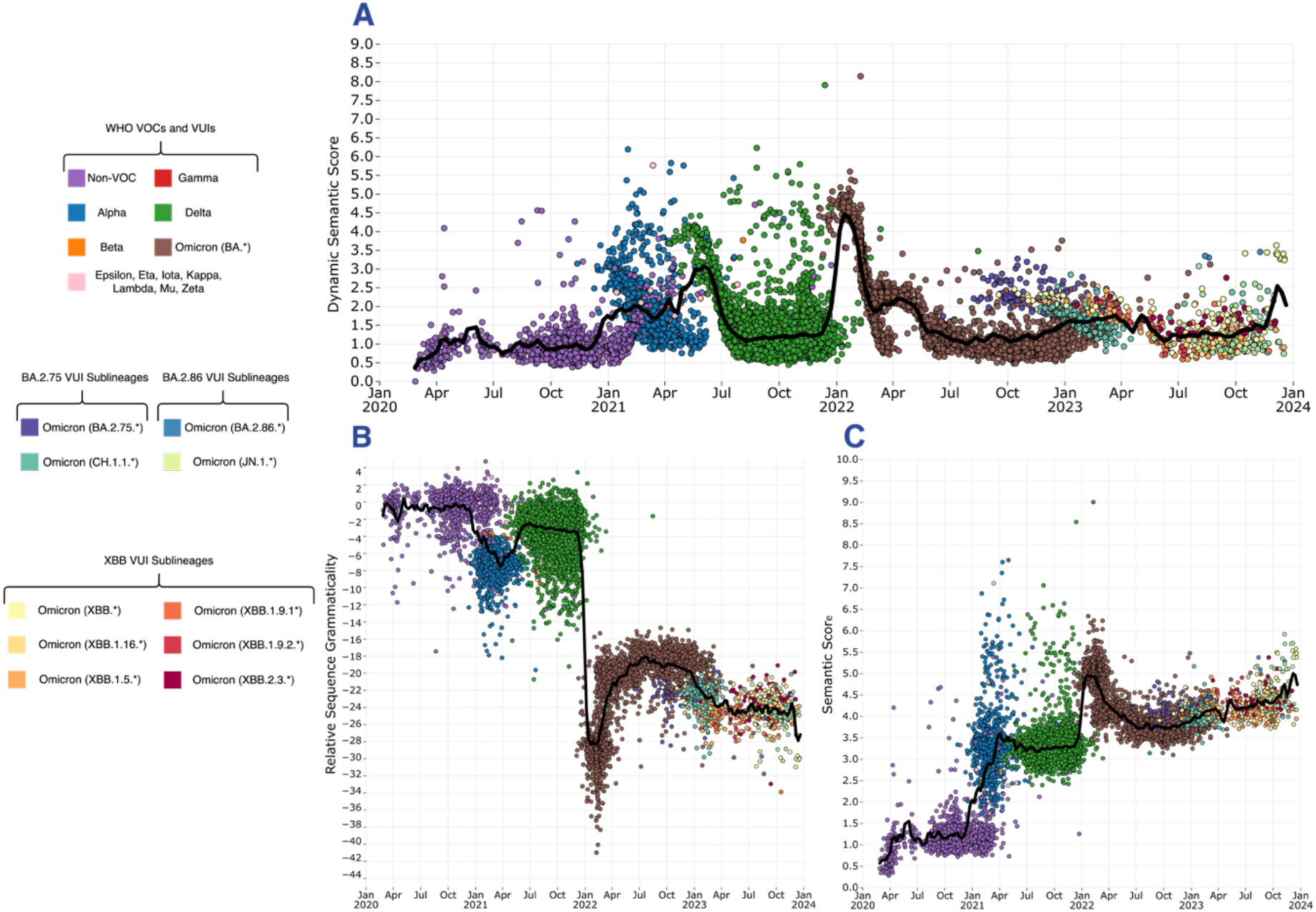
(A) UK SARS-CoV-2 spike sequences through the pandemic. Each point represents a sequence cluster with 99.9% sequence similarity. Dynamic semantic scores were calculated for each sequence cluster, with the black line showing the mean sliding score. (B) Relative sequence grammaticalities for each of the haplotype spikes. (C) Semantic scores for each of the haplotype spikes.

Some Delta sequences have higher relative sequence grammaticalities than the reference SARS-CoV-2 sequence (Figure 6B), which suggests Delta may more “fit” than other SARS-CoV-2 sequences when all likelihoods are considered. The metrics also highlight outlier sequences, although these might be related to factors beyond sequence properties such as epidemiology which make variant prediction difficult for both experimental and computational methods. Dynamic embedding references can assist with modelling the shifting immune landscapes SARS-CoV-2 is evolving within. Emergent antigenically distinct variants are likely to cause repeated infection by SARS-CoV-2^40^. Consequences of this are quickly becoming apparent through the necessity for updated vaccines, with recent variants evading previously neutralising antibodies^41,42^. We capture major VOC transitions but also incremental variants like JN.1. The heavily mutated BA.2.86 lineage acquired an L455S substitution^43^ to produce JN.1, which subsequently became the dominant lineage circulating. Its identification by the dynamic reference score highlights the approach as a potentially viable horizon scanning method that does not simply count mutations, but accurately models them.

In conclusion, we have shown that the PLM metrics grammaticality and semantic score reveal characteristic properties of the SARS-CoV-2 spike protein sequence. Using just a single spike sequence we were able to determine the likely regions of variability in the spike protein, and identify regions where mutations were most likely to impact the structure and function of the protein. We describe how PLMs can map out evolutionary landscapes, identify epistatic effects and provide a method for horizon scanning of viral sequences of interest. Unlike other methods for predicting potential SARS-CoV-2 variant success, although representing improvements, for example, Ito et al.^5^ and Thadani et al.^4^, these need protein structures and multiple sequence alignments for training. In our implementation, use of the pre-existing model ESM-2, negates the need for training and permits characterisation based on a single sequence. While there are clear areas for further improvement, this represents a paradigm shift for interpreting variation within protein sequences and could be applied to a novel pathogen when data is extremely sparse.

While ESM-2 is a good predictor of the computational and experimental metrics used to understand how mutations impact protein structures, these metrics do not correlate strongly with any one feature, indicating that PLM-derived scores are capturing something novel about protein sequences. PLMs are now being used for DMS and variant effect prediction tasks across an impressive array of datasets, and often outcompete other methods^44^. They are helping improve the effectiveness of antibodies^45^ and even generate new proteins^46^. Our results here affirm that these models are useful and can be applied to great effect to understand novel viral pathogens. PLMs, thus, offer an exciting glimpse into the future of language modelling within biology, and we should continue to press on with understanding the possibilities as well as the limitations of what these models can do.

## Methods

### ESM-2 and in silico DMS

SARS-CoV-2 spike proteins were acquired by filtering the GISAID database (https://gisaid.org) for the earliest sequences from each PANGO lineage with a fully intact spike protein sequence. The sequence was then embedded in the ESM-2 model to produce an embedding for the sequence. For the DMS data and for the embedding scores, the ESM-2 three billion parameter variant was used. The semantic score is equivalent to the L1 (Manhattan) distance between the embedding of the reference sequence (the spike protein from the original Wuhan-Hu-1 SARS-CoV-2 genome) and each of the PANGO spike proteins. The grammaticality of a sequence is calculated as the product of the probabilities of each amino acid at each position in the spike protein. The probabilities come from a softmax of the last layer of the embedding and range between 0 and 1. However, many of the probabilities are small and for numerical stability the probabilities are represented in the log space. Relative grammaticality is the same as the grammaticality, except the probability of a reference sequence is subtracted from the probability of the variant sequence so that the score is relative, in this case, to the SARS-CoV-2 reference sequence Wuhan-Hu-1. The sequence grammaticality represents the summed log-likelihoods of every reference position in the sequence, rather than just the mutated positions. For the DMS computations, a sequence was produced for every potential amino acid at every position in the spike protein. Each sequence was then embedded to calculate semantic scores and relative grammaticalities for every mutation.

### Evo-velocity

We used the evo-velocity package^37^ to embed the initial sequences using the ESM-2 650M parameter variant, and then performed velocity analysis and spearman’s rank for the sample dates against the pseudo-time. The smaller model was used primarily due to hardware limitations of using the larger model, although in a number of cases this model has been show to equal or even do better than the larger model.

### Epistasis Experiments

The epistasis experiments used a SARS-CoV-2 BA.1 spike protein sequence and embedding this using the ESM-2 three billion parameter variant. Due to the EPE insertion, the logits for this position were subsequently removed to map logits to spike’s three-dimensional protein structure. The BA.1 sequence contains several mutations relative to the Wuhan-Hu-1 SARS-CoV-2 reference spike sequence. Each of these mutations was reverted one by one back to the reference position, and the likelihood differences upon reversion were recorded. To eliminate noise, changes less than two standard deviations from the mean across all reverted mutations were removed and deemed not significant.

### Dynamic embeddings and horizon scanning

Dynamic embeddings were computed by first gathering UK GISAID data and clustering sequences into variants with 99.9% similarity. These haplotype variant clusters produced 11,272 sample date labelled haplotypes with a sequence returned for each cluster. Each haplotype embedding was measured against a mean of the embeddings from the prior three-month period using the L1 distance, i.e., the semantic score. For sequence grammaticalities, mean sequence grammaticalities were calculated in a similar way and differences measured.

### Assessing embeddings scores with known metrics

The language model’s metrics were first assessed using a Spearman’s rank against several known biological scores. Next, Support Vector Regression (SVR) was used as a simple model to fit the model scores as well as the embeddings and logits to the biologically relevant metrics. Models were fit to the data using five-fold cross validation, and a linear kernel for the SVR. Model results were reported as the average spearman’s rank between the folds, with the error bars as +/-1 standard deviation from the mean. To assess statistical divergence between metrics at every site scores were normalised to between 0 and 1, preserving rank order. The Jensen Shannon distance for each comparison was then calculated for every site independently using the philentropy (v.0.8.0) package in R. Amino acid positions used to define protein regions were as follows: S1 N-terminal (14-306), S1 RBD (331-528), S1 C-terminal (529-686), S2 (687-1273), all other positions were classed as indeterminate. Greater JSD implies greater divergence between two distributions. Significance was determined by Mann Whitney U test with Bonferroni correction.

### Selection analysis signals and entropy

Signals of ancestral evolutionary selection in the animal (bat and pangolin) sarbecovirus most closely related to SARS-CoV-2 referred to as the “nCoV” clade (see Lytras et al.^47^) were inferred on a set of 167 sarbecovirus genomes, accounting for recombination by inferring selection separately in each non-recombinant segment. These results are published in Martin et al.^48^ and presented in more detail in the following Observable notebook: https://observablehq.com/@spond/ncos-evolution-nov-2021. Sites under negative selection were inferred using the FEL and sites under positive selection using MEME^49^ by testing on internal branches of the nCoV clade. Sites denoted as conserved have the same amino-acid residue among all sarbecovirus sequences in the analysis. The variability of each site in the SARS-CoV-2 sequence was obtained from the entropy of the predicted distribution of credible evolutionary states.

### Antibody accessibility and substitution probabilities

Structure-based epitope score, referred to as “accessibility”, which approximates antibody accessibility for each spike protein amino acid position, was calculated using BEpro software^50^ for a Woo *et al.* model of the Wuhan spike. Scores relating to substitution probabilities, namely, ESST probability, Log PSSM and predicted ΔΔG, were obtained for every possible single amino acid substitution for the 6VXX SARS-CoV-2 spike structure (note that only values for Chain A are included in the results as data is generally identical across all three chains). ESST probability values were calculated using Environment Specific Substitution Table (ESST)^51^ after local structural environments were calculated by JOY^35^ Log Likelihood substitution values were calculated using Position Specific Scoring Matrices (PSSM) with the DELTA-BLAST^52^ algorithm in BlastX. It should be noted that this is a sequence-based method, so residue numbering does not match the numbering in 6VXX and values are available for residues not described by the 6VXX PDB file. ΔΔG values were predicted by FoldX5^53^ software. FoldX uses empirical energy functions to predict the energetic effect of mutations to protein stability. The predicted ΔΔG quantifies the change in the free energy of unfolding between the wild-type and mutated structure. The 6VXX structure was first repaired with the *RepairPDB* function to fix residues with bad torsion angles, van der Walls’ clashes or total energy. Substitutions were then performed using the repaired structures on all three chains simultaneously using the *BuildModel* function, giving the change in free energy of unfolding, with negative values implying stabilising mutations.

### Deep mutational scanning data

The receptor binding deep mutational scanning data was taken from experimental DMS studies of the SARS-CoV-2 from Yisimayi *et al.* (2024) and Starr *et al.* (2022)^33,34^. The Wuhan-Hu-1 scores were taken as is, while the variant average score was calculated by averaging the scores for each position between each of the SARS-CoV-2 variant specific DMS results^34^. For the RBD mutational escape values we utilised the high-throughput mutation antibody escape profiling results presented in Yisimayi *et al.* (2024)^33^ This study used a panel of 1,350 monoclonal antibodies against all possible RBD substitutions. The backbone virus used was the SARS-CoV-2 BA.5 variant instead of Wuhan-Hu-1, however this should still provide the most comprehensive dataset of unique substitutions’ effect on antibody escape. Our mutational escape metric is the average of the raw antibody escape values for each substitution on each site of the RBD across all tested monoclonal antibodies. The full spike DMS data was taken from Dadonaite *et al.* (2023) and DMS scores (“*Binding”*, “Escape” and “Entry”) are from a BA.2 and XBB.1.5 spike backbone and averaged together to give the presented score^32^.

## Data Availability

The initial sequences for each PANGO lineage in Figure 5 were extracted from GISAID, their accession numbers can be found here: https://doi.org/10.55876/gis8.240620pm. The SARS-CoV-2 spike sequences used for Figure 6 are from GISAID, their accession numbers can be found here: https://doi.org/10.55876/gis8.240621ma. The code as well as data can be found on GitHub: https://github.com/kieran12lamb/PLM_SARS-CoV-2.

## Acknowledgements

We gratefully acknowledge all data contributors, i.e., the Authors and their Originating laboratories responsible for obtaining the specimens, and their Submitting laboratories for generating the genetic sequence and metadata and sharing via the GISAID Initiative, on which this research is based. Thanks for the neural network icon made by Becris from www.flaticon.com. The authors acknowledge funding from the UK Medical Research Council (MRC, MC_UU_12014/12, MC_UU_00034/5, MR/V01157X/1 and a Doctoral Training Programme in Precision Medicine studentship for KDL, MR/N013166/1), the Wellcome Trust (220977/Z/20/Z), the UK Research and Innovation (UKRI) to the G2P-UK consortium (MR/W005611/1) and G2P2 consortium (MR/Y004205), and the COVID-19 Genomics UK Consortium (COG-UK), which was supported by funding from the MRC, part of UKRI, the UK National Institute of Health and Care Research (MC_PC_19027) and Genome Research Limited, operating as the Wellcome Sanger Institute. For the purpose of open access, the author has applied a Creative Commons Attribution (CC BY) licence to any Author Accepted Manuscript version arising.

## Supplementary figures

**Supplementary Figure 1.**
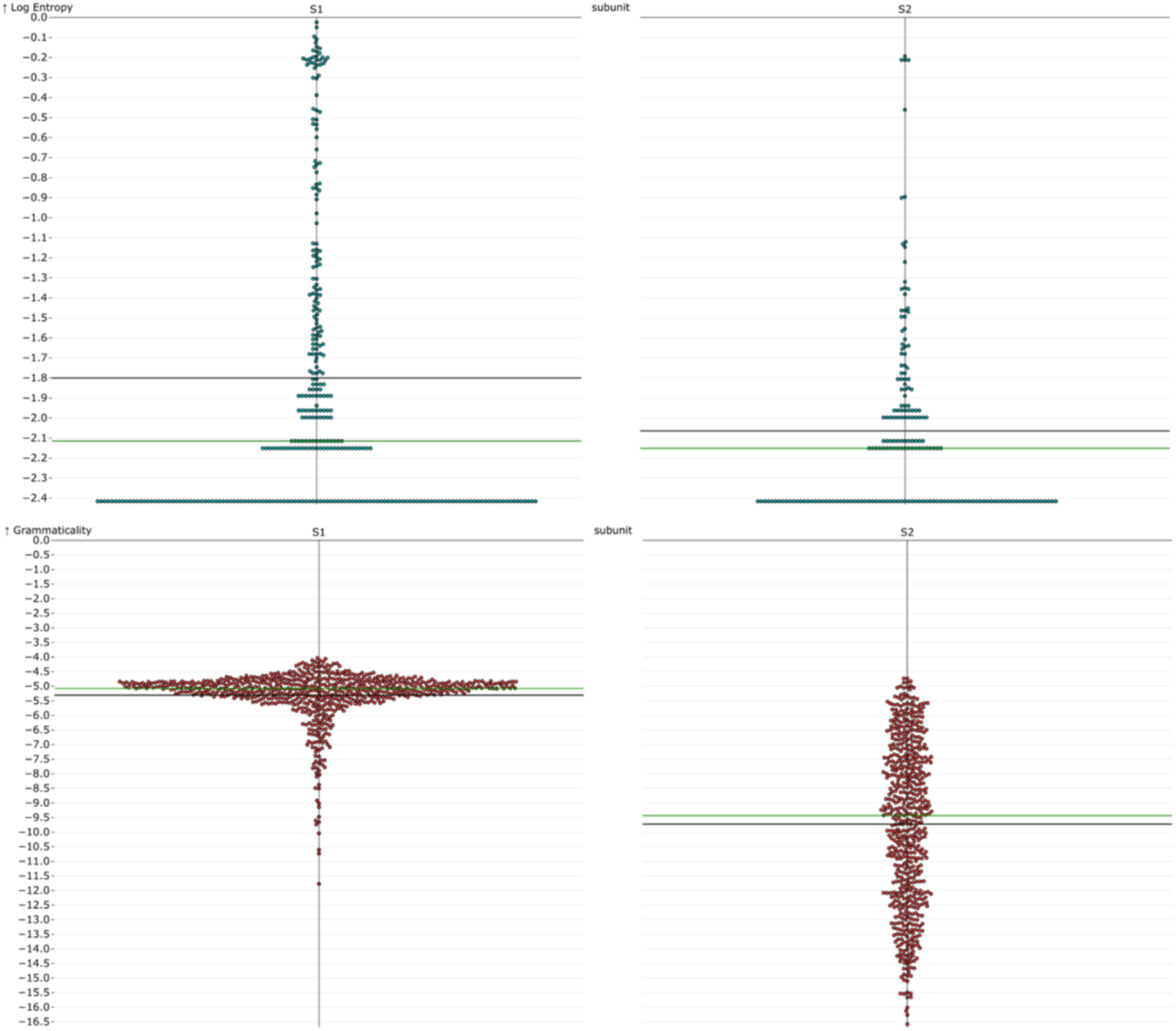
Swarm plots for the distributions of entropy and grammaticality for each of the SARS-CoV-2 spike protein subunits S1 and S2. The black line shows the mean while the green shows the median. For both entropy and grammaticality, the S2 subunit has on average lower scores compared to the S1.

**Supplementary Figure 2.**
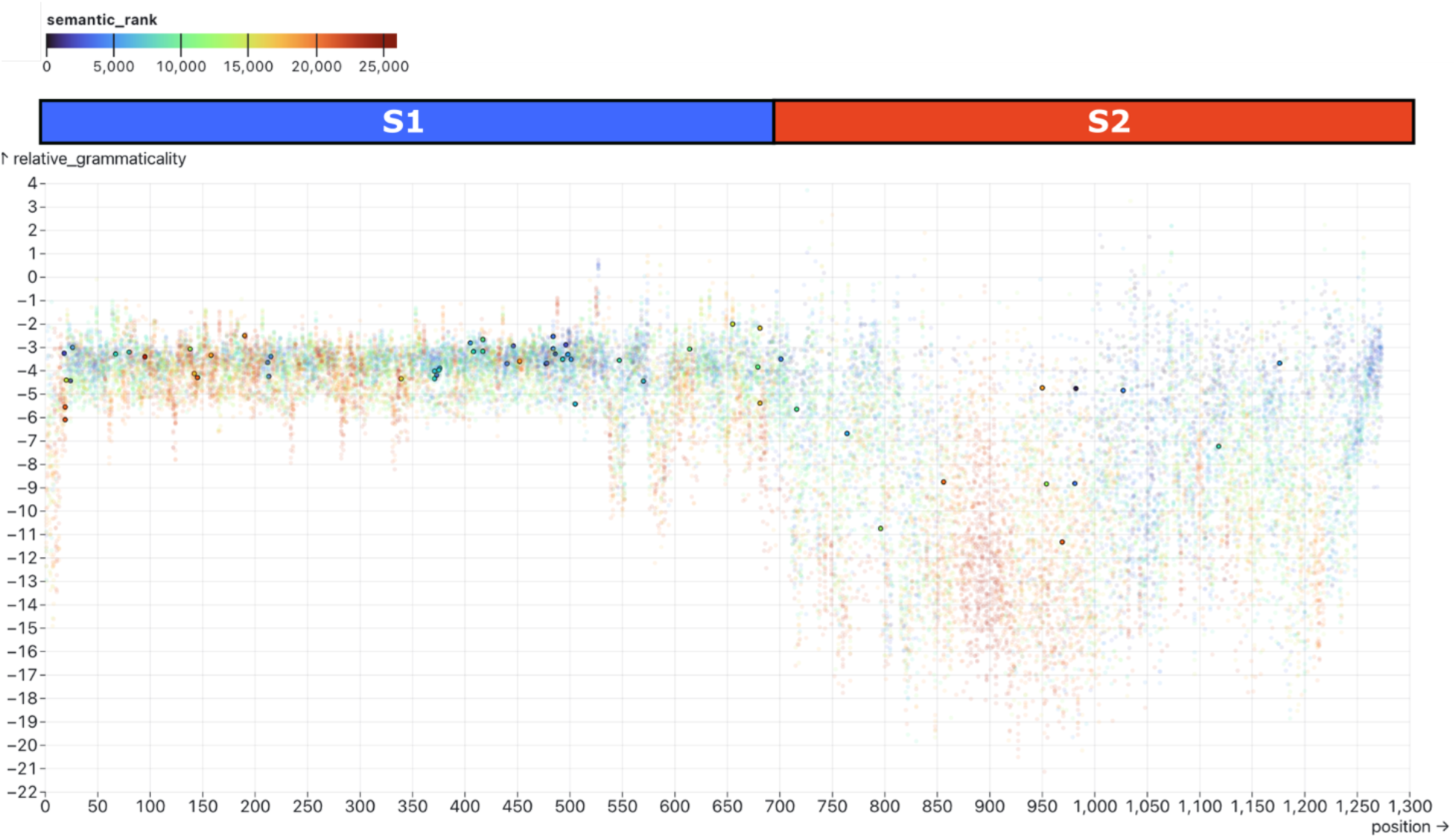
DMS plot with only the substitutions observed in SARS-CoV-2 VOC sequences highlighted. Most of the mutations occur in the spike protein S1 region which the model predicts has a higher likelihood of mutations, i.e., a higher grammaticality.

**Supplementary Figure 3.**
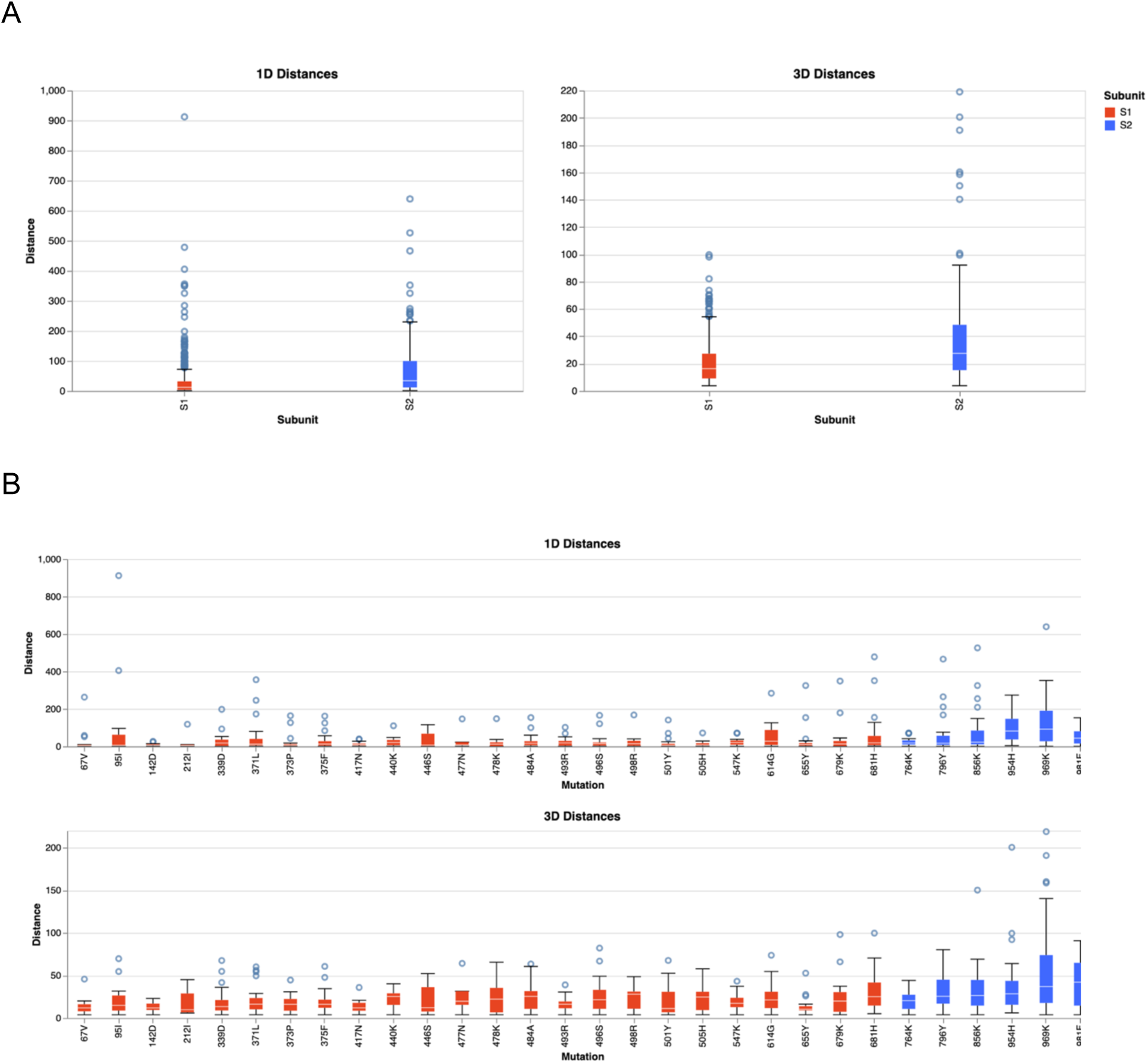
(A) Boxplots showing the distribution of distances between mutations and the positions affected by mutations in each subunit. (B) Boxplots showing the distribution of distances between mutations and the positions for each mutation.

**Supplementary Figure 4.**
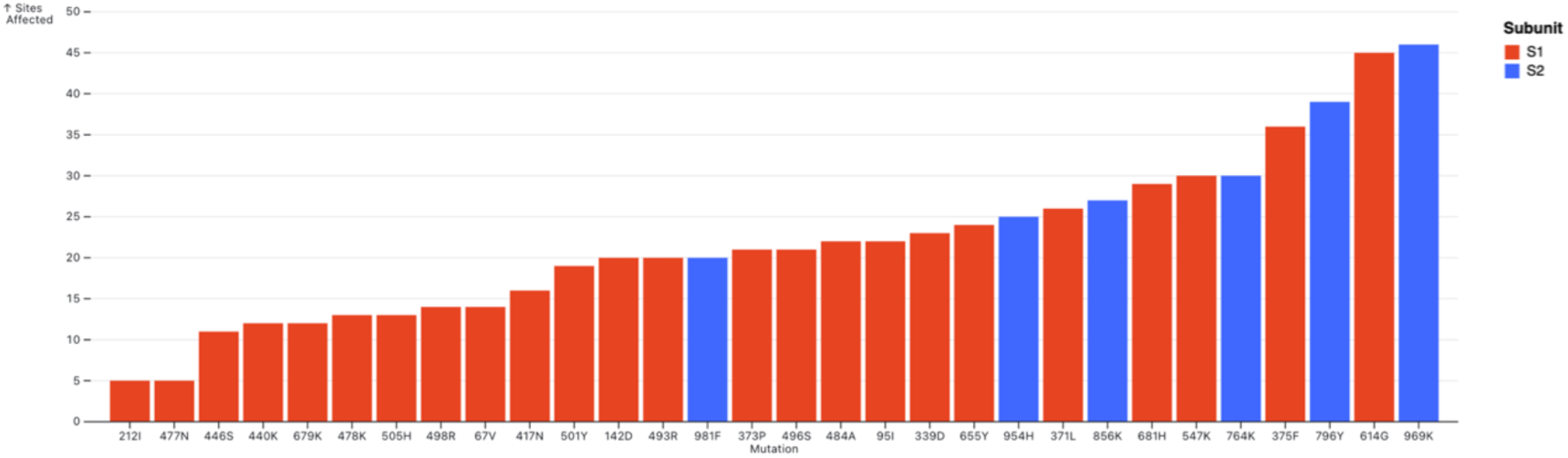
Number of sites with a significant (+/-2 deviations from mean) change in probability for each SARS-CoV-2 BA.1 reversion change.

**Supplementary Figure 5.**
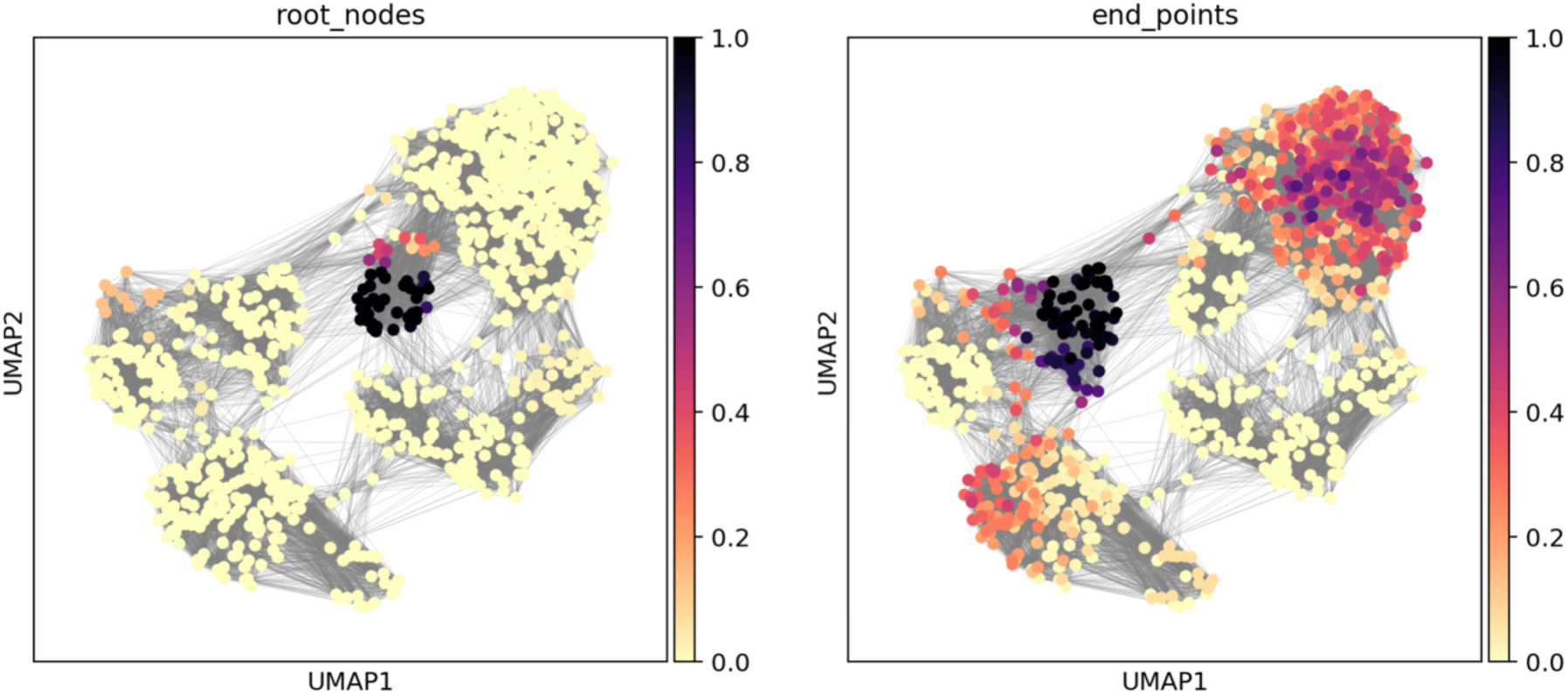
Predicted root nodes and endpoints identified by running Markov diffusion process over the weighted edges of the evo-velocity network. The root nodes are correctly identified as the Sarbecovirus spike sequences, with Omicron VOC sequences predicted as the end nodes.

**Supplementary Figure 6:**
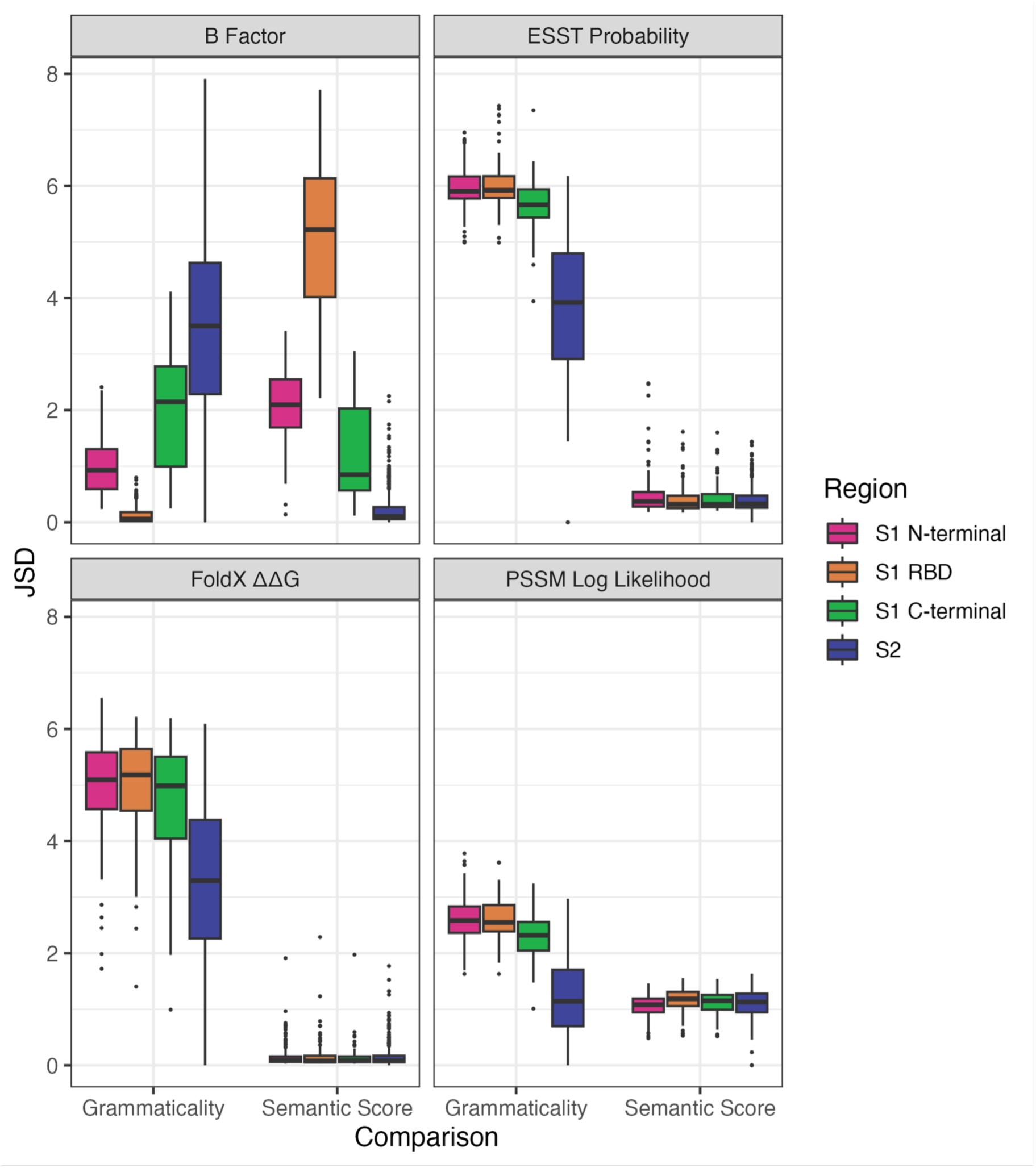
Boxplots showing Jensen Shannon distances (JSDs) between semantic score or grammaticality and other computational metrics. JSD was calculated for every site, colours show the distribution of scores in different protein regions. Comparisons between S1 regions and S2 are all significant by Mann Whitney U test following Bonferroni correction (p<0.05) except for the following comparisons: FoldX ΔΔG vs semantic score; all S1 vs S2 comparisons, Log Likelihood vs semantic score; S1 C-terminal vs S2, ESST probability vs semantic score; S1 RBD vs S2, S1 C-terminal vs S2.

**Supplementary Figure 7.**
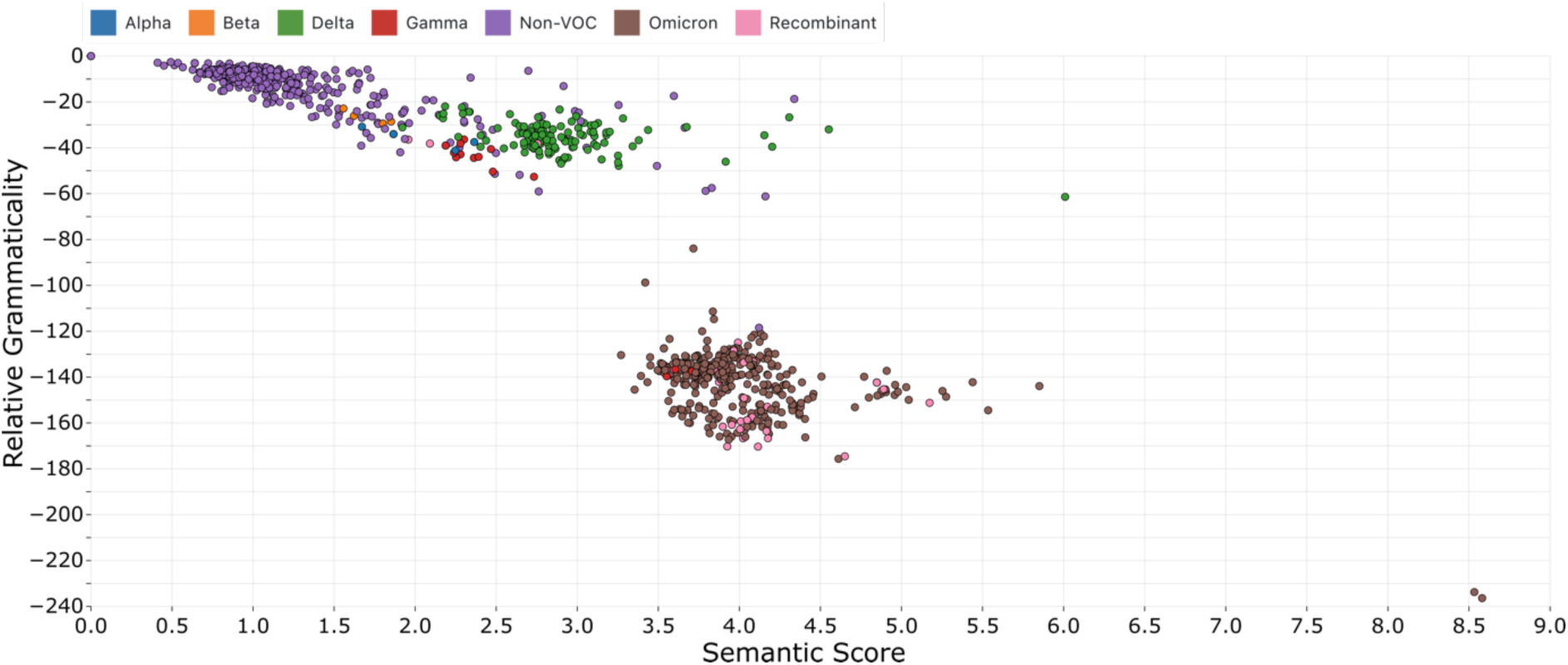
SARS-CoV-2 Pango lineage representative sequences plotted by their semantic scores and relative grammaticalities.

**Supplementary Figure 8.**
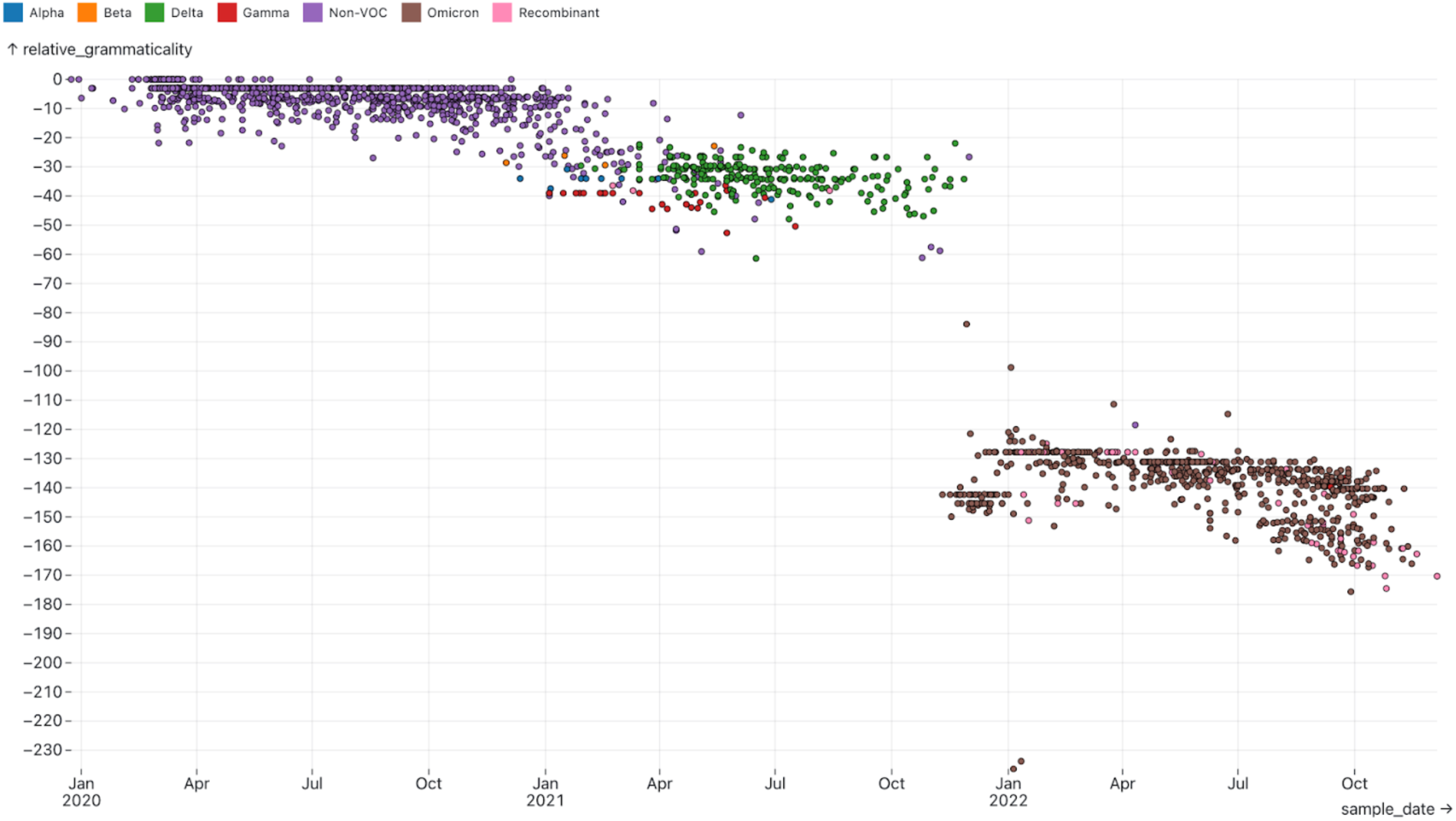
SARS-CoV-2 Pango lineage representative sequences plotted by their relative grammaticalities against sampling date.

